# Performance evaluation of adaptive introgression classification methods

**DOI:** 10.1101/2024.06.12.598278

**Authors:** Jules Romieu, Ghislain Camarata, Pierre-André Crochet, Miguel de Navascués, Raphaël Leblois, François Rousset

## Abstract

Introgression, the incorporation of foreign variants through hybridization and repeated backcross, is increasingly being studied for its potential evolutionary consequences, one of which is adaptive introgression (AI). In recent years, several statistical methods have been proposed for the detection of loci that have undergone adaptive introgression. Most of these methods have been tested and developed to infer the presence of Neanderthal or Denisovan AI in humans. Currently, the behaviour of these methods when faced with genomic datasets from evolutionary scenarios other than the human lineage remains unknown. This study therefore focuses on testing the performance of the methods using test data sets simulated under various evolutionary scenarios inspired by the evolutionary history of human, wall lizard (*Podarcis*) and bear (*Ursus*) lineages. These lineages were chosen to represent different combinations of divergence and migration times. We study the impact of these parameters, as well as migration rate, population size, selection coefficient and presence of recombination hotspots, on the performance of three methods (VolcanoFinder, Genomatnn and MaLAdapt) and a standalone summary statistic (Q95(*w*, *y*)). Furthermore, the hitchhiking effect of an adaptively introgressed mutation can have a strong impact on the flanking regions, and therefore on the discrimination between the genomic windows classes (*i.e.* AI/non-AI). For this reason, three different types of non-AI windows are taken into account in our analyses: independently simulated neutral introgression windows, windows adjacent to the window under AI, and windows coming from a second neutral chromosome unlinked to the chromosome under AI. Our results highlight the importance of taking into account adjacent windows in the training data in order to correctly identify the window with the mutation under AI. Finally, our tests show that methods based on Q95 seem to be the most efficient for an exploratory study of AI.

## Introduction

Hybridisation is the reproduction of individuals belonging to differentiated gene pools. As long as reproductive isolation remains incomplete, hybridisation and backcrossing can lead to introgression, *i.e.* the incorporation of genetic material from a donor population into the genome of a recipient population (Anderson and Hubricht, 1938). Mallet (2005) estimated that hybridisation occurs in over 25% of plant species and 10% of animal species, and more evidence of interspecific hybridisation and introgression has emerged since then in plants and animals following the advent of high-throughput sequencing technologies (Taylor and Larson, 2019; Twyford and Ennos, 2012). Indeed, the barriers between species are not completely impermeable, allowing genetically distinct species to be maintained over time in the presence of gene flow (Harrison and Larson, 2014; Suarez-Gonzalez et al., 2018). While interspecific introgression is increasingly recognised as an important process in the evolution of biodiversity, its prevalence (in terms of the proportion of species involved) and importance (in terms of the proportion of the genome that is affected) remain poorly understood (Edelman and Mallet, 2021).

Hybridization and introgression have many potential evolutionary outcomes. They can lead ultimately to the accumulation of deleterious mutations (Adavoudi and Pilot, 2022), or to the loss of genetic diversity by the merging of the two parent species (Runemark et al., 2019), or even to the extinction of the parent species through genetic swamping (Todesco et al., 2016). Introgression can also allow the acquisition of alleles that increase the fitness of carrier individuals, a process known as adaptive introgression (AI; Burke and Arnold, 2001; Kim and Rieseberg, 1999). Although there are few examples to date, adaptive introgression is thought to occur in both animals and plants (*e.g.* Burgarella et al., 2019; Hedrick, 2013). In some cases, adaptive introgression could even play an important role in the speciation process as proposed in Darwin’s finches (Grant and Grant, 2019) or butterflies of the genus *Heliconius* (Pardo-Diaz et al., 2012; Rosser et al., 2024).

Proving the presence of adaptive introgression in a species is challenging, because evidence is needed to support that the genomic regions involved come from another species and either that they provide a selective advantage on the fitness of the individuals carrying them or that they exhibit signs of positive selection in the recipient species (Burgarella et al., 2019; Suarez-Gonzalez et al., 2018). The first studies of adaptive introgression looked independently for introgression and for selection signals in genetic sequences. However, the signals left by adaptive introgression in genetic sequences can be quite distinct from signals left by neutral introgression or by selective sweeps of non-introgressed variants (Burgarella et al., 2019; Racimo et al., 2015; Setter et al., 2020; Taylor and Larson, 2019). For this reason, classification methods have been developed to detect genomic signatures specific to adaptive introgression (Gower et al., 2021; Racimo et al., 2017; Setter et al., 2020; Zhang et al., 2023). These methods aim at identifying genetic patterns specific to adaptive introgression, such as summary statistics focusing on excess of allele sharing between the introgressed population and the introgressing population (Racimo et al., 2017), genomic scans identifying patterns of diversity specific to adaptive introgression (Setter et al., 2020) or machine learning algorithms trained on data simulated under adaptive introgression scenarios (Gower et al., 2021; Zhang et al., 2023).

Most of these methods have been developed and tested on human genetic data to identify potentially adaptively introgressed regions caused by ancient hybridization among groups of humans: *Homo sapiens*, *Homo neanderthalensis* and Denisovans (human evolutionary models reviewed in Racimo et al., 2015). In addition, methods using machine learning algorithms have been trained only on genetic datasets simulated under human scenarios. The behaviour and performance of these methods on organisms with evolutionary scenarios distant from the human lineage are therefore unknown. Using a simulation-based inference method trained on a particular biological model to infer a biological process on another model could degrade the method’s performances. The signals left by adaptive introgression depend on life history traits and evolutionary history of the biological model. The history of connections between populations, their sizes and divergence times, as well as the availability for sampling of introgressing, introgressed and non-introgressed populations, and the strength of selection are features specific to the biological model and the study, which influence the signature of AI in the genome and therefore the performance of the methods. In the case of non-model species, the knowledge of demographic history and genomic parameters such as mutation and recombination rates is often limited. For these species, it is difficult to choose the scenario under which genetic data should be simulated to train machine learning methods to detect AI, and more generally to define a score threshold differentiating regions under AI from other regions. Indeed, most current studies trying to detect AI in non-model species use VolcanoFinder (Setter et al., 2020), the only method that does not require the definition of an evolutionary model or data of the introgressing species (Liu et al., 2022; Pawar et al., 2023; Wang et al., 2023) to be applied on a given data set. However, some authors have suggested that alternative simulation-based methods are robust to misspecification of demographic history (Gower et al., 2021; Zhang et al., 2023). It is thus important to assess the robustness of these methods on genetic data generated by evolutionary histories not explored in the training set.

Furthermore, as these methods have been published recently, no study has compared their performance since the original articles. One aim of our work is therefore to compare the performance of published adaptive introgression detection methods in relation to evolutionary parameters possibly affecting this detection: migration time and rate, strength of selection on the introgressed allele, effective population size, recombination rate, and divergence time between taxa. To do so, we simulated genetic datasets under a diversity of evolutionary scenarios inspired by the life history traits of three case studies: humans, wall lizards (*Podarcis*) and bears (*Ursus*). We set the human demographic scenario as our reference, and used it to evaluate the impact of variation in migration rate and effective population sizes as well as heterogenous recombination rate on classification performance. The *Podarcis hispanicus* complex includes at least eight species that inhabit the Iberian Peninsula where they are sympatric or parapatric, meeting in narrow contact zones (Pinho et al., 2009; Renoult et al., 2009). Various levels of genomic and mitochondrial introgression have been identified between several species (Caeiro-Dias et al., 2021; Pinho et al., 2009; Renoult et al., 2009; Yang et al., 2021). Their divergence times are on the order of a million generations (Kaliontzopoulou et al., 2011), much older than divergence between humans and neanderthals dating from less than 20,000 generations (Prüfer et al., 2014). Scenarios based on the *Podarcis* evolutionary history thus allow us to examine the impact of the divergence time between donor and recipient populations. Human migration times are relatively recent (∼2,000 generations) and a similar timing of gene flow has been proposed for *Podarcis* (gene flow was made possible by the post-glacial expansion of populations, Pinho et al., 2011). prompting us to consider an additional scenario with older migration. Introgression between polar bears (*Ursus maritimus*) and brown bears (*Ursus arctos*) occurred between 10,000 and 32,000 generations ago (Lan et al., 2022; Liu et al., 2014). We thus devised a third scenario inspired by this case to evaluate the effect of migration time on the detection of AI.

In this study, we perform simulation tests of the performance of four different existing AI classification methods: VolcanoFinder (Setter et al., 2020), MaLAdapt (Zhang et al., 2023) and Genomatnn (Gower et al., 2021) and the summary statistic Q95(*w*, *y*) (Racimo et al., 2017). Each method has been developed, trained (for simulation-based methods), and evaluated for a different type of non-AI windows (*i.e.* genomic windows that do not carry AI alleles, see below). Gower et al. (2021) uses neutral introgression windows simulated independently of AI windows, while Zhang et al. (2023) uses windows that do not carry the mutation under AI but share the same demographic and adaptive history (*i.e.* generated by the same simulation), and that can be physically linked to the AI-windows or not. Methods that do not require training, such as VolcanoFinder, also use different non-AI window classes for tests.

As the aim of all these classification methods is to specifically identify windows carrying introgressed advantageous mutations, the types of non-AI windows used in the train set to achieve this objective is extremely important. Testing for the presence of AI in a genetic dataset composed of windows adjacent to the AI window using a method not trained with this type of window may result in the identification of false positives, because hitchhiking can lead to the identification of false AI signals in such adjacent windows. Therefore, we also explore the impact of intra- and inter-chromosomal hitchhiking on performance, by taking into account in the test sets adjacent windows linked to the window carrying the mutation under AI, as well as a second chromosome harbouring only neutral windows but potentially affected by hitchhiking. Indeed, although hitchhiking is mainly considered as the selective effect of an advantageous allele on genetic variation at adjacent neutral alleles linked to it (Maynard Smith and Haigh, 1974), which we call intra-chromosomal hitchhiking in this article, its effect extends to other chromosomes, an effect named here inter-chromosomal hitchhiking. This hitchhiking follows from any initial statistical association between a selected allele and alleles at physically unlinked loci, in particular the associations created by the immigration event as well as their persistence in the first generations following immigration.

We thus evaluate the performance of four AI classification methods to detect genomic regions containing an allele under various scenarios of AI inspired by demographic models of the human, wall lizard and bear lineages, to illustrate the impact of varying a parameter of interest, and compared how the four tested methods are robust to different types of non-AI windows. Our results highlight the differences among the methods, and provide a guideline on what method to use for the study of adaptive introgression. Our simulation tests suggest that the simple genome scan method based on one of the Racimo et al. (2017) summary statistics generally has better performance in the classification of genomic regions under adaptive introgression than all other more complex methods in most of the scenarios considered.

## Material and methods

### Adaptive introgression model

We simulate genetic data under an adaptive introgression model where a beneficial allele is transferred by gene flow from a donor population to a recipient population. The full model is composed of five populations (Fig. 1): an outgroup population (O), a recipient population (R), a non-introgressed sister population of the recipient one (RS), a donor population (D), and a non-introgressing sister population of the donor one (DS). The donor’s sister population is used in our study as a stand-in for the donor population to mimic empirical situations when the latter is unknown or where genomic data are not available from it (Setter et al., 2020). Genetic data sampled from different combinations of these populations are used by the evaluated methods.

**Figure 1.**
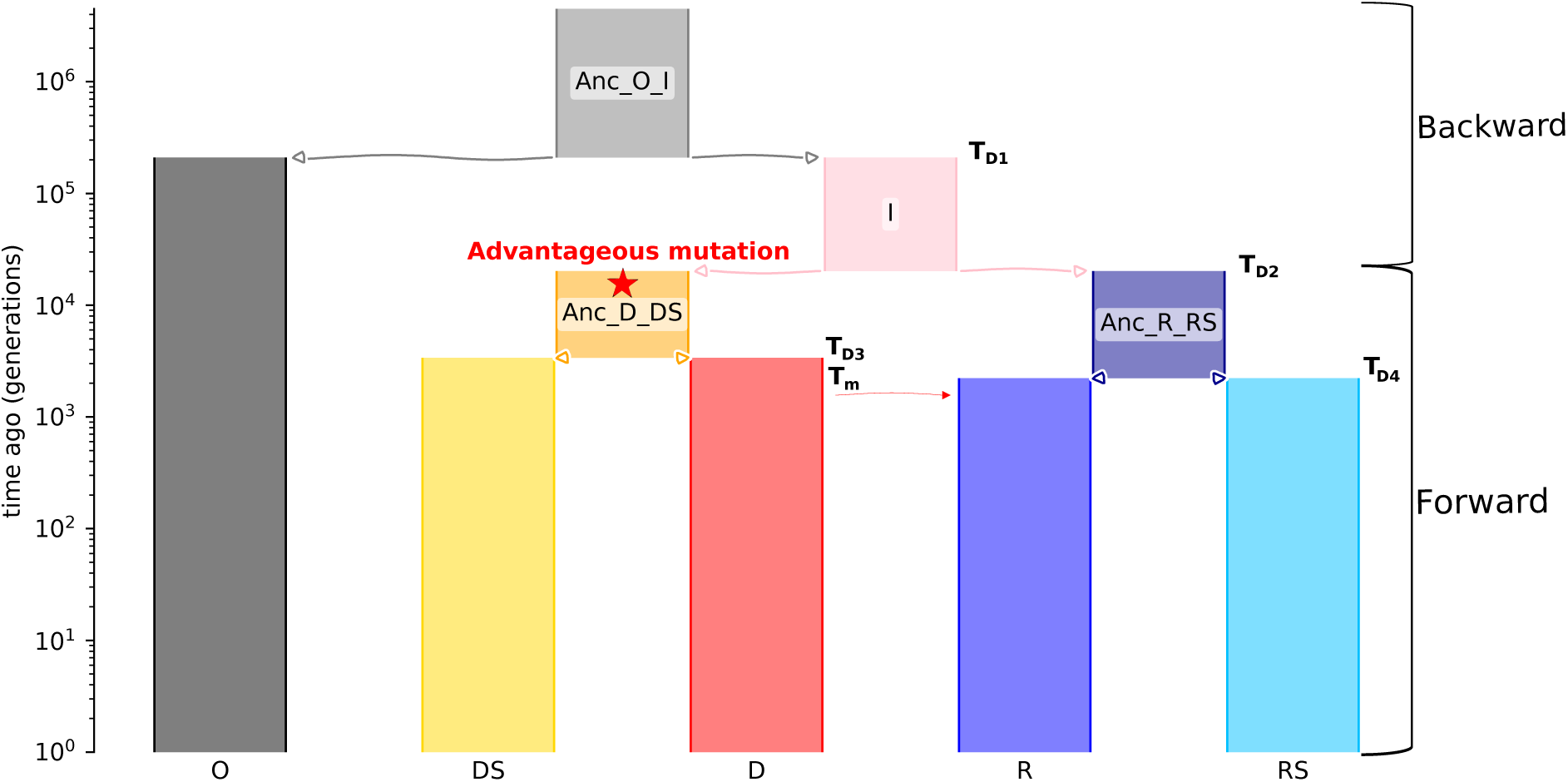
Demographic model used in this study. **Anc_O_I** = Outgroup and Ingroup ancestral population, **O** = Outgroup population, **I** = Ingroup population, **Anc_D_DS** = Donor and non-introgressing ancestral population, **Anc_R_RS** = Recipient and non-introgressed ancestral population. **DS** = sister to the donor population (non-introgressing population), **D** = Donor population (introgressing population), **R** = Recipient population (introgressed population), **RS** = Recipient sister population (non-recipient population), **T_D1_** = Divergence time between **O** and **I**, **T_D2_** = Divergence time between **Anc_D_DS** and **Anc_R_RS**, **T_D3_** = Divergence time between **D**, **DS**, **T_D4_** = Divergence time between **R** and **RS** and, **T_m_**= migration generation time. The arrows and triangles with a white background represent divergence events between populations, the red arrow represents the migration event.

Population sizes are constant throughout the duration of the simulation and fixed to a single value *N* for all populations. At the end of the simulation, a given number of individuals is sampled in each population, except the donor one. Except for reference scenarios of neutral introgression (NI) for which we consider a single chromosome, we simulate diploid genomes with two chromosomes, of length 1Mb each, where neutral mutations can occur at a mutation rate *µ* and (intra-chromosomic) recombination events at a rate *r*, both rates being expressed per base per gamete (*i.e.* per generation). The different chromosomes are simulated by setting a recombination rate of 0.5 between the two genomic regions defining them. To simulate an adaptive introgression event, a single beneficial mutation (with selection coefficient *s*) is positioned at the centre of one of the two haploid genomes of a single individual within the ancestral donor population (Anc_D_DS in Fig. 1). The beneficial mutation is then potentially transferred from the donor population to the recipient population during a single generation of migration, with an immigration rate *m*. The selection coefficient for the advantageous mutation does not change over time or between populations (*i.e. s* is the same for Anc_D_DS, D, DS and R). Simulation are discarded when the selected mutation is lost in AI scenarios, or when no genetic material is introgressed in NI scenarios.

Demographic parameters values are chosen to produce demographic scenarios inspired by the evolutionary history of the human, *Podarcis* or *Ursus* lineages (Table 1). Variable parameter values among all the scenarios are shown in Table 2. Each demographic scenario is used to test the impact on performance of varying a specific parameter. The Human demographic model is used to define the reference scenario (*i.e.* scenario with which the performances of the other scenarios are compared), whose event times are inspired by the “PapuansOutOfAfrica_10J19” model from the library for population genetic simulation, stdpopsim (Adrion et al., 2020). In this scenario, *Homo neanderthalensis* and *Homo sapiens* are the donor and recipient species, respectively, and *Pan troglodytes* the outgroup. For the reference scenario, *µ* is set to 1.2×10*^−^*^8^ mutation per base per gamete, the recombination rate is set to the same value as the mutation rate; *m* and *N* are set to *m* = 10*^−^*^2^ and 10, 000 respectively, and *s* is set to 10*^−^*^2^, to simulate intermediate selection. In other human scenarios, the values of a single parameter are modified, in order to test the impact of the variation of this specific parameter on performance. The other human scenarios are therefore built to test the impact of variation in immigration rate (*m* = 10*^−^*^1^ and 10*^−^*^3^), small effective population sizes (*N* = 200) and the presence of recombination hotspots. Hotspots are simulated as areas with high recombination rates (*r* = 5 × 10*^−^*^7^) over a 1kb span, occurring within each 100kb window (similarly to observed patterns reviewed by Myers et al., 2006). Outside these hotspots, the recombination rate is lower (*r* = 4 × 10*^−^*^9^), maintaining the average recombination rate to 1.2 × 10*^−^*^8^. In the first hotspot scenario, the hotspot located in the window under AI is positioned 20kb away from the mutation under selection, while in the second scenario, the mutation is situated at the centre of the hotspot.

**Table 1.**
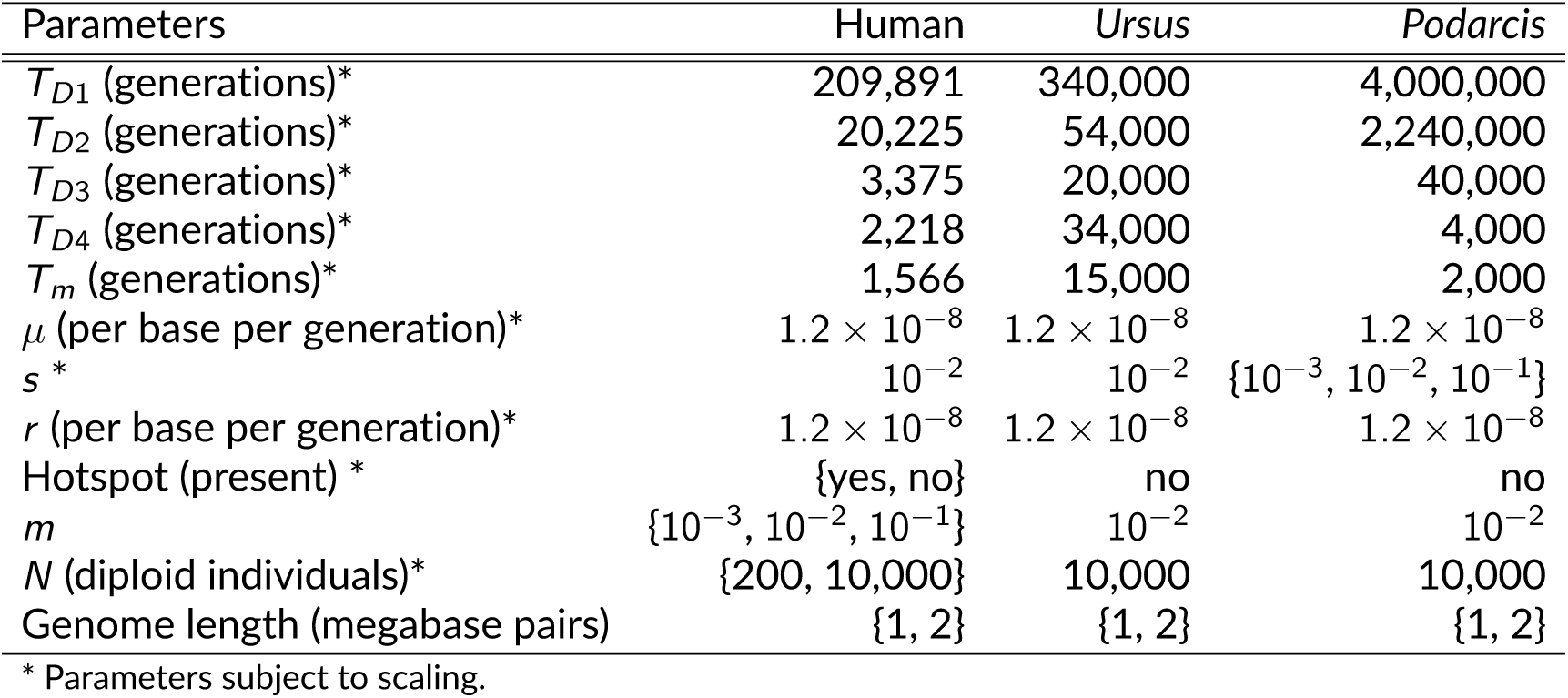
Parameter values used in the demographic scenarios.

**Table 2.**
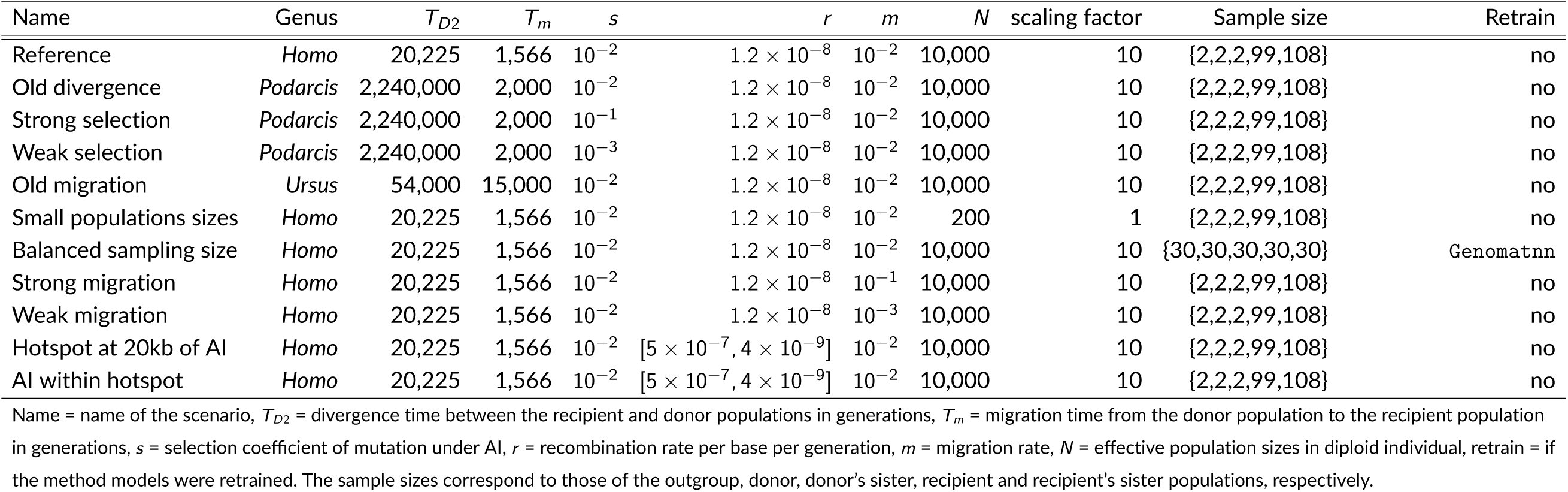
Variable parameter values for each scenario.

The *Podarcis* scenarios are designed to test the impact of an ancient divergence between the D and R populations (*T_D_*_2_), inspired by the potentially adaptive introgression found between the species *Podarcis carbonelli* (D) and *Podarcis bocagei* (R; Caeiro-Dias et al., 2021, 2023; Gaczorek et al., 2023; Pinho et al., 2009), which are thought to have diverged 2.24 million generations ago (5.6 million years ago, Kaliontzopoulou et al., 2011). In this scenario, *Podarcis muralis* was chosen as the outgroup species. The *Podarcis* scenarios also made it possible to assess the impact of selection strength, test sets are generated with different selection coefficients representing strong, intermediate and weak selection scenarios (*s* values of 10*^−^*^1^, 10*^−^*^2^ and 10*^−^*^3^).

Finally, the scenario based on bear demographic events shows the impact of an earlier migration time than that used for the human scenarios. The genus *Ursus* was chosen because an introgression event between the polar bear and the brown bear occurring between 10,000 and 32,000 generations ago (100,000 and 320,000 years ago) has previously been inferred (Lan et al., 2022; Liu et al., 2014). In this scenario, the American black bear (*Ursus americanus*) is the outgroup species, the polar bear (*Ursus maritimus*) the donor species and the brown bear the recipient species (*Ursus arctos*). For both *Podarcis* and *Ursus* scenarios, the values of *m*, *N*, *µ* and *r* are identical to the reference human scenario, to allow the comparison of results.

We use a combination of forward and backward (coalescent) simulators to generate the genetic data. First, msprime (Baumdicker et al., 2022; Kelleher et al., 2016) simulates backward in time the ancestral recombination graph (ARG) of the whole population from the divergence of populations ancestral to the donor and recipient populations (*i.e.*, at time *T_D_*_2_ in Fig. 1), until the most recent common ancestor. SLiM (Haller and Messer, 2019) is then used to simulate, forward in time and starting from msprime’s ARG, the more recent section of the evolutionary scenario which includes selection, *i.e.* from *T_D_*_2_ until present. SLiM’s ARG of the individuals sampled in current populations is used by msprime to add neutral mutations to generate the sample polymorphism.

For most scenarios, sample sizes are set to 99, 108, 2 and 2 for the recipient population, the non-introgressed population, the sister population of the donor, and the outgroup population, respectively, corresponding to the values used by the tested simulation-based classification methods (Genomatnn and MaLAdapt, Gower et al., 2021; Zhang et al., 2023). However, these two methods used samples from the donor population for their train and test set where we used samples from the sister population of the donor. In order to test the impact of sample size, a test set was simulated under the reference human scenario, with balanced samples of 30 individuals per population. Haplotypic and genotypic matrices for all samples are then obtained from the ARG with mutations, using the tskit (Ralph et al., 2020) and scikit-allel packages (Miles et al., 2024). As forward simulations can be highly time-consuming, a scaling factor of 10 was applied to all parameter values (except *m*, since migration occurs as a one-time event limited to a single generation in which the same proportion of the population is replaced by migrants in the original and scaled models) to drastically reduce simulation times: *s*, *µ* and *r* are multiplied by the scaling factor, and the times of events and *N* are divided by the scaling factor (as in Gower et al., 2021; Hoggart et al., 2007; Zhang et al., 2023). Although the use of a scaling factor on selection coefficient and population size can bias the forward simulation results (Dabi and Schrider, 2024), its use in our analysis is justified not only by the fact that simulating the test sets under the *Podarcis* scenario without it would be extremely time-consuming (*i.e.* over 100 times longer than with the scaling factor, representing hundreds of thousands of CPU hours), but also because the simulation-based classification methods (Genomatnn and MaLAdapt) used it to generate their training set. Table S1 compares the parameter values used to train the simulation classification methods (MaLAdapt and Genomatnn) and our test sets.

### Methods to detect adaptive introgression

The performance of four adaptive introgression classification methods: VolcanoFinder (Setter et al., 2020), MaLAdapt (Zhang et al., 2023) and Genomatnn (Gower et al., 2021) and a method based on the summary statistic Q95(*w*, *y*) (Racimo et al., 2017) are evaluated in this work. All of them are classification methods whose objective is to detect regions of the genome that harbour a beneficial mutation originating from the donor population (AI region).

The Q95(*w*, *y*) statistic (Racimo et al., 2017) is one of the first summary statistics developed to specifically capture the signal of adaptive introgression in genetic data. This summary statistic can be calculated for both phased and unphased genotypes. It gives the 95% quantile of the distribution of derived allele frequencies in the recipient population, for alleles that have a frequency equal to or higher than *y* in the donor population and lower than *w* in the sister population of the recipient population. This summary statistic (named Q95 thereafter), with *w* = 0.01 and *y* = 1, and computed on 50kb non-overlapping windows, is used here as a score value to detect adaptive introgression.

VolcanoFinder (Setter et al., 2020) is the first adaptive introgression classification method to have been developed. This method aims to identify adaptive introgressed alleles recently fixed in the recipient population, considering only the genetic information from the introgressed population. It uses an optional outgroup population to polarize the alleles and does not require phased haplotypes. For a given genomic position, VolcanoFinder calculates a composite likelihood ratio for the site frequency spectrum (SFS) around that focal site, comparing an AI model and a reference SFS acting as empirical null model. In our tests, we used the SFS of the whole chromosome as reference. To compute the composite likelihoods, VolcanoFinder can estimate the strength of selection and the genetic differentiation between the recipient and donor populations or use user-specified fixed values. We choose to let VolcanoFinder infer them, as recommended in Setter et al. (2020). We set the interval between each tested site to 1000bp. The scores per position given by the method (*i.e.* the composite likelihood ratio) are further converted into a single score per 50kb window, corresponding to the highest score observed in the window considered. To obtain a relative score (*i.e.* between 0 and 1 as for the other methods), the score of each window is then divided by the maximum score observed among all windows and all analysed datasets for a given scenario.

Genomatnn (Gower et al., 2021) is a simulation-based classification method that uses a deep learning algorithm (convolutional neural network, CNN), trained on 100kb windows simulated independently under different scenarios with samples from the donor, recipient and recipient’s sister populations, to detect adaptive introgression. Genomatnn gives a probability for each 100kb tested window to have been affected by an adaptive introgression event and can use both phased and unphased data. The training set includes sequences simulated under adaptive introgression, neutral introgression and classical selective sweep scenarios, and is used to classify genomic regions (windows) as AI or non-AI. We used two pre-trained CNNs (Nea_to_CEU_af-0.25_2918410235.hdf5 and Nea_to_CEU_af-0.05_2250018620.hdf5) described in the article presenting the method and corresponding to CNNs trained on simulations filtered so that the frequency of the beneficial allele in the recipient population was greater than or equal to 5% and 25% (respectively Genomatnn_0.05 and Genomatnn_0.25 thereafter; Gower et al., 2021). We use these originally trained CNNs to test their robustness when using test datasets simulated under a different demographic scenario. We also re-trained Genomatnn with balanced sample sizes of 30 individuals per population without changing other parameters for the test of the effect of sample sizes. For this, 100,000 datasets were simulated under each AI, neutral introgression and classical selective sweep processes with the Genomatnn pipeline.

MaLAdapt (Zhang et al., 2023) is also a simulation-based classification method and uses a machine learning (ML) algorithm (extra tree classifier, Geurts et al., 2006), trained on simulations of a 5Mb chromosome phased genome under different scenarios with samples from the donor, recipient and recipient’s sister populations, to classify genomic regions (50kb windows) as AI or non-AI. For each window, MaLAdapt computes a score corresponding to the probability that the window has experienced an AI event. Contrary to Genomatnn, MaLAdapt is not trained on raw sequences, but on a set of summary statistics calculated on each window. These summary statistics were chosen by Zhang et al. (2023) to capture patterns of genetic diversity, linkage disequilibrium and population differentiation, that are informative about introgression, selection and adaptive introgression. Contrarily to Zhang et al. (2023), we use non-overlapping windows and do not consider the presence of exons in the genome. The number of exons per window, a summary statistic used by Zhang et al. (2023), has been removed because this information is only available for annotated genome, which is rarely the case for non-model species. Furthermore, its use rests on simulations assuming that selection only takes place in exons, which is not consistent with evidence of selection outside exons (Jo and Choi, 2015). This assumption should systematically lead to the classification of windows without exon as non-AI windows and thus artificially improve the method’s performance. We also do not compute Kelly (1997)’s *Z_nS_*summary statistic, because it is computationally intensive to calculate and because *Z_nS_* does not carry pertinent information according to the classification importance score of the extra-tree classifier algorithm of MaLAdapt (Zhang et al., 2023). Re-using the original simulated training set from Zhang et al. (2023), we retrained two extra-tree classifiers without the two statistics mentioned above, one with the Q95(0.01, 1.0) statistic and the other without (respectively MaL-Adapt_Q95 and MaLAdapt_noQ95 thereafter), which are equivalent to models 4 and 6 from Zhang et al. (2023) but without *Z_nS_* and number of exons. Summary statistics from train and test sets are standardised using their mean and variance computed on the training set (whereas Zhang et al., 2023, used the mean and variance computed on the analysed test set).

Inferences were thus carried out on non-overlapping 50kb windows (for VolcanoFinder MaL-Adapt and the method based on Q95) or 100kb windows (for Genomatnn), and using phased genotypes of the donor’s sister population, recipient and recipient’s sister populations for Genomatnn and MaLAdapt or from the recipient population only for VolcanoFinder.

### Method performance evaluation

To identify candidate AI-windows, the four methods select windows whose score is higher than a given threshold. The threshold values need to be defined for each data set, as they depend on the sampling scheme and evolutionary history of the samples. Here, we varied the score threshold value used to classify the windows as AI or non-AI and computed true and false positive rates to generate receiver operating characteristic (ROC) curves for the different types of non-AI windows and calculate the area under the ROC curve (AUC, Bradley, 1997). The use of these evaluation metrics (ROC and AUC) allows us to avoid defining score thresholds for each method that would only be relevant for a given scenario and sampling scheme.

For all methods tested, we define a region being under adaptive introgression as a window containing the advantageous mutation originating from the donor population (named AI window). But the definition of the “reference” non-adaptive introgression (non-AI) regions (*i.e.* the null model) differs for the different classification methods: (i) VolcanoFinder does not assume a specific null model in terms of evolutionary history but rather defines an empirical null model from the genome-wide SFS of the sampled individuals in the recipient population; (ii) Genomatnn uses a train set including, as non-AI windows, neutrally introgressed (NI) windows and nonintrogressed windows with a beneficial site (classic sweep) with a class ratio of 1:1:1 (AI:NI:classic sweep), simulated independently of each other; and (iii) MaLAdapt uses (1) all windows of the 5Mb chromosome other than the one carrying the advantageous mutation (referred to here as adjacent windows) as non-AI windows, so that AI and adjacent non-AI windows are jointly simulated and can be physically linked, (2) selective sweep windows simulated independently of the AI windows. The AI:non-AI class ratio of the final training set is 1:2. By including adjacent windows as non-AI windows in its training sets, MaLAdapt treats as non-AI some windows that have undergone hitchhiking and therefore potentially do not have the same genetic diversity as neutral introgression windows simulated independently of the AI window. Indeed, depending on the strength of selection, recombination and demographic events, the hitchhiking effect of the advantageous mutation can have a strong impact on the evolution of the frequency and distribution of linked neutral alleles. In the context of adaptive introgression analysis, hitchhiking certainly has an effect on inference and may notably lead, for example, to the detection of false AI signals on windows adjacent to the beneficial allele. Furthermore, the effects of hitchhiking may extend to other chromosomes than the one carrying the mutation, as even free recombination (*r* = 1*/*2) might not immediately suppress the inter-chromosomal linkage disequilibrium in the first generations after the introgression of the beneficial haplotype. None of the AI classification methods tested here take this inter-chromosomal hitchhiking into account in the definition of its training set nor their sensitivity with respect to such hitchhiking in the (simulated) data has been tested.

To examine the effect of different forms of hitchhiking on the distribution of scores computed by each method, we included neutral windows affected by intra- or inter-chromosomal hitchhiking in the test sets. The train sets were not modified in this respect relative to the original definitions of the methods (the definition of the Genomatnn train set ignores hitchhiking; for MaLAdapt, it includes intra-chromosomal hitchhiking; the two other methods are not machinelearning methods with a train-set concept). The effect of inter-chromosomal hitchhiking is assessed by simulating genomes containing two chromosomes of 1Mb each, the first one carrying the introgressed adaptive mutation and the second one being fully neutral, and by comparing scores for windows from the second chromosome to scores within the first one. The effect of intra-chromosomal hitchhiking is assessed by examining the variation in distribution of scores for windows at different distances from the adaptive mutation on a 1Mb chromosome. Last, we also generated test sets of a 1Mb chromosome simulated without any selection (so containing only neutral introgression windows) to represent Genomatnn types of neutral non-AI windows.

For each scenario explored, we simulated one test set of 200 simulated data. We represent the distributions of scores over the 200 replicates as function of the distance to the beneficial mutation when relevant (*i.e.* not for the chromosome 2 and independent neutral chromosome, where hitchhiking affects the test sets).

## Results

We compare the performance of the four methods described above, initially in our human reference case and then in relation to seven factors of variation: the strength of selection (*s*), the migration rate (*m*), the population size (*N*), the recombination rate heterogeneity (*r*), the divergence time between the D and the R population (*T_D_*_2_) and the migration time (*T_m_*). Those comparisons are reported for different types of non-AI windows: windows simulated independently under a neutral introgression scenario (NI), neutral windows not physically linked to the beneficial mutation (*i.e.* on the second chromosome, chro2), or neutral windows on the same chromosome as the beneficial mutation (*i.e.* adjacent windows). The consideration of those three different classes of non-AI window classes allows us to evaluate false positive rates (FPR) in different regions of the genome and in the case of complete absence of AI. ROC curves (Fig. 2 and S1 for adjacent windows, Fig. 3 and S2 for independent neutral introgression windows and Fig. S3 for second chromosome windows) and AUC (Table S3 to S5) express the expected performance of each method and ease the comparisons between them. On the other hand, plots of scores along the chromosomes are used to explore in more details the effects of hitchhiking on their performance (Fig. 4 and Fig. 5, and Fig. S4 to Fig. S11).

**Figure 2.**
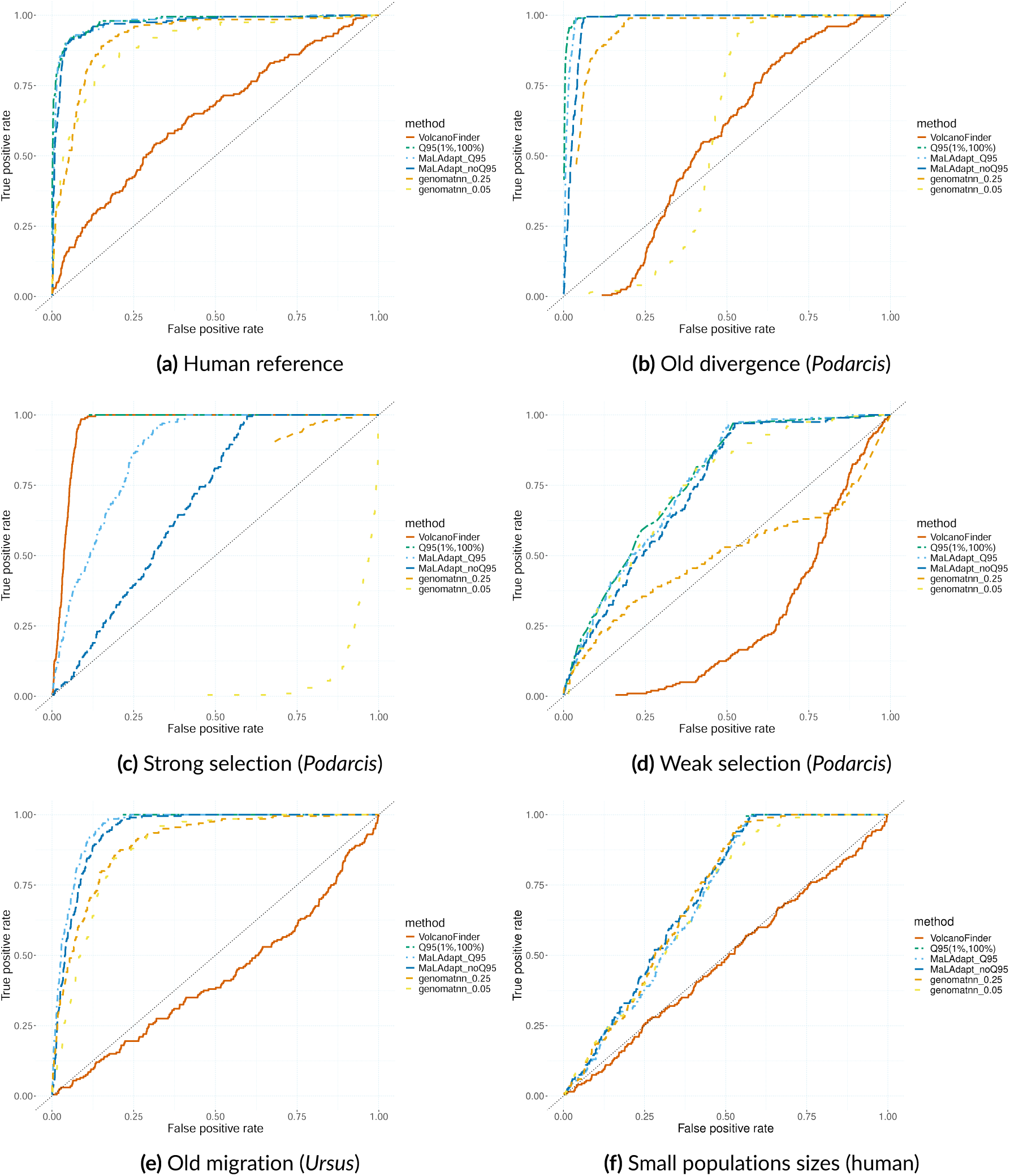
ROC curve for adaptive introgression classification methods for 200 test sets simulated under human, *Podarcis* and *Ursus* scenarios with a chromosome containing a mutation under various selection strength and adjacent windows as non-AI windows. True positive window = window with AI mutation, False positive windows = adjacent windows.

**Figure 3.**
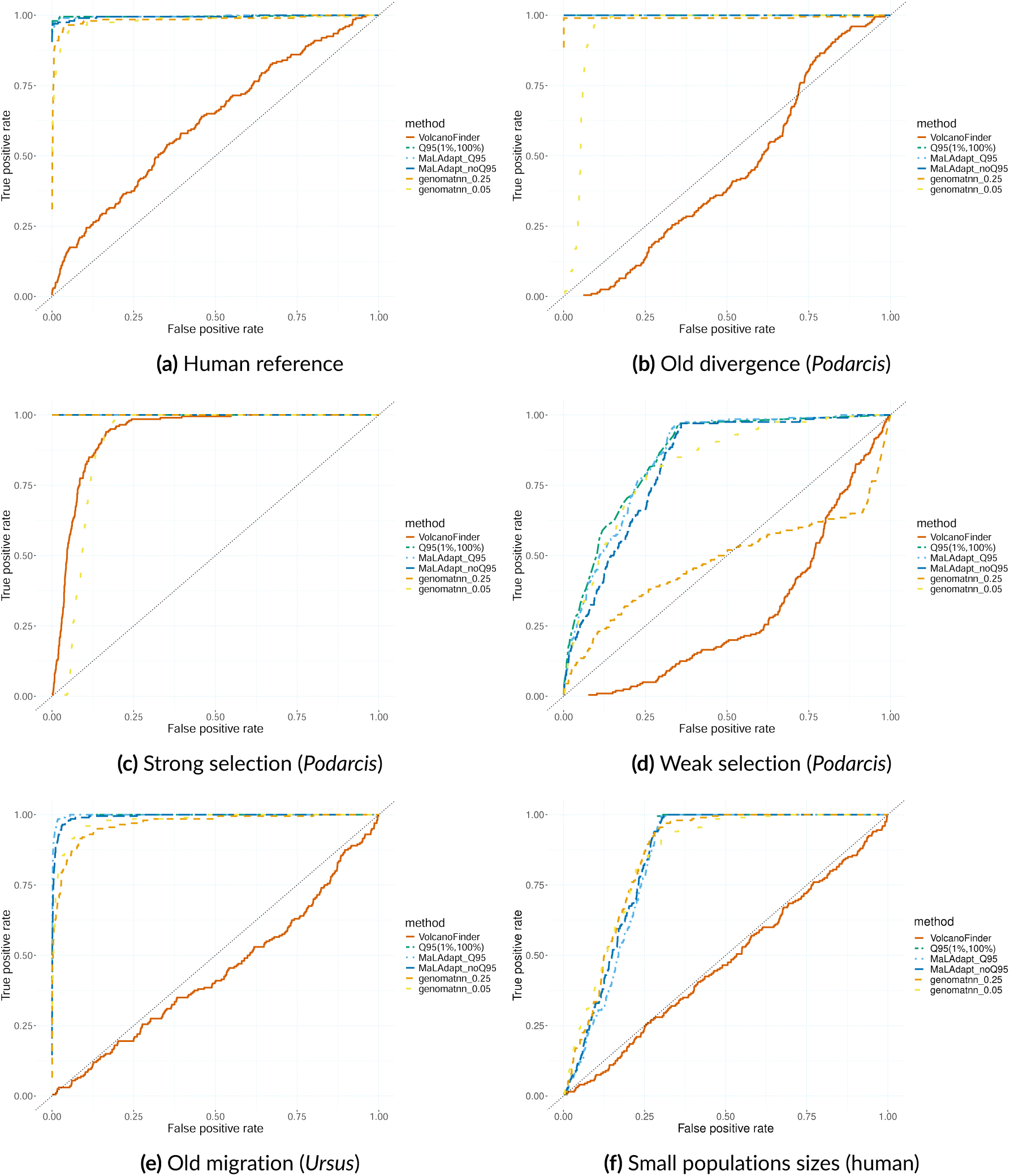
ROC curve for adaptive introgression classification methods for 200 test sets simulated under human, *Podarcis* and *Ursus* scenarios with a chromosome containing a mutation under various selection strength and independent neutral introgression windows as non-AI windows. True positive window = window with AI mutation, False positive windows = Independent neutral introgression (NI).

**Figure 4.**
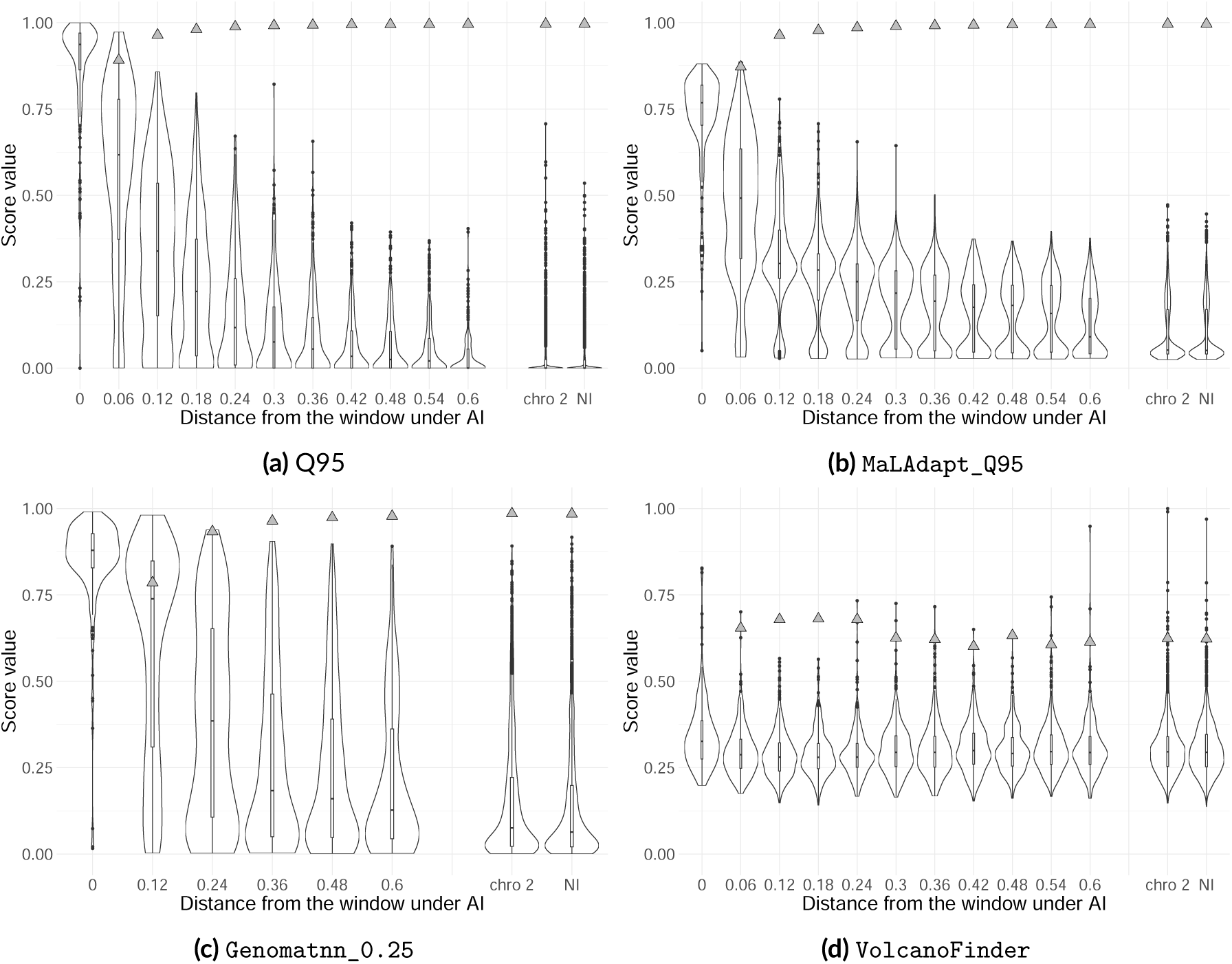
Score values of 200 test sets for the different windows considered under human reference scenario with intermediate selection strength (*s* = 10*^−^*^2^). Scores from the first chromosome are represented as function of the distance (cM) of the windows to the site under selection (0 indicates the window containing it), followed by scores from the second neutral chromosome (“chro 2”) and from an independently simulated neutral chromosome (NI) for reference. The areas under the ROC curves (AUC) for each test set containing AI windows and each class of non-AI windows are represented by the grey triangles. Each violin plot is from at least 200 windows.

**Figure 5.**
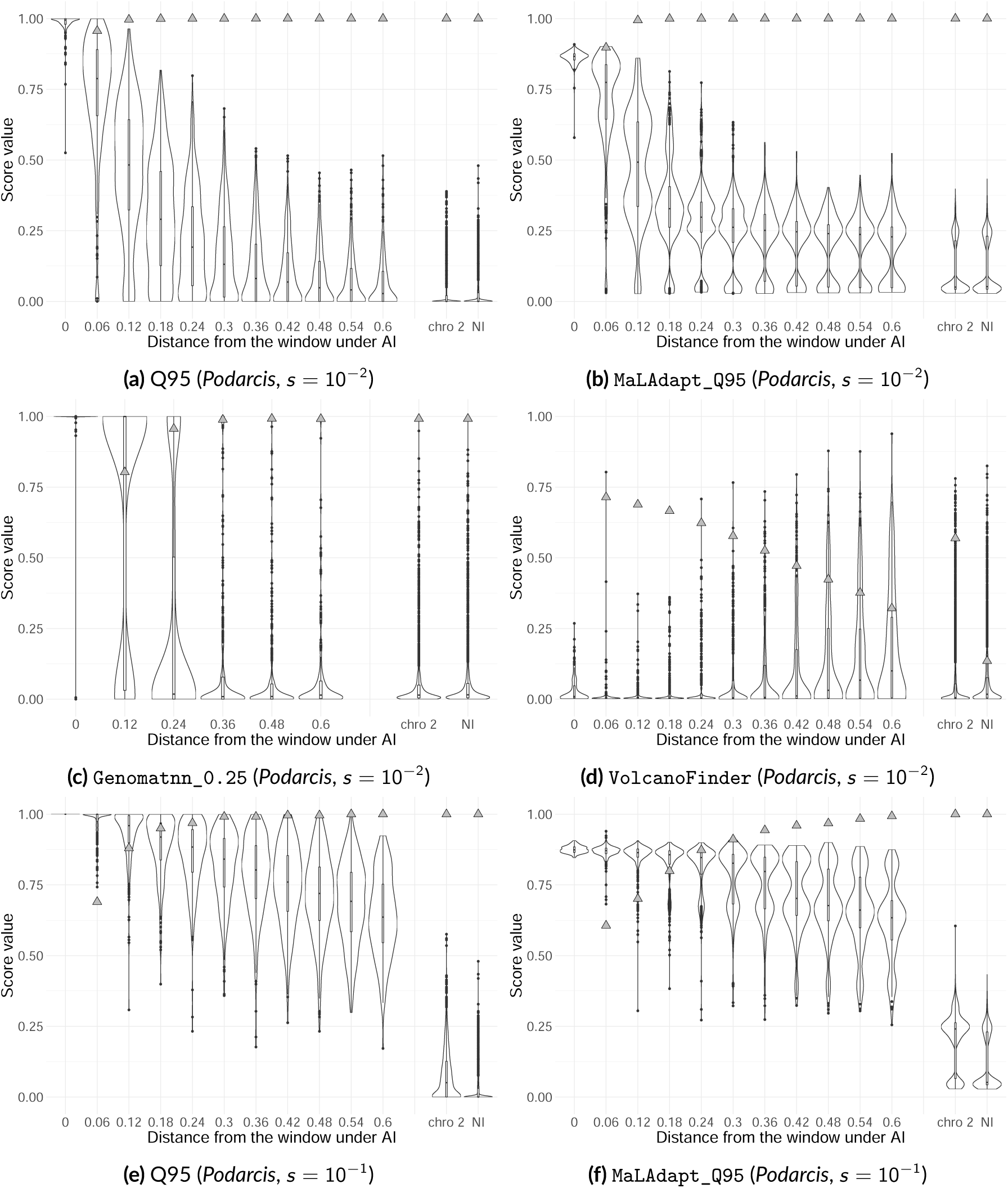
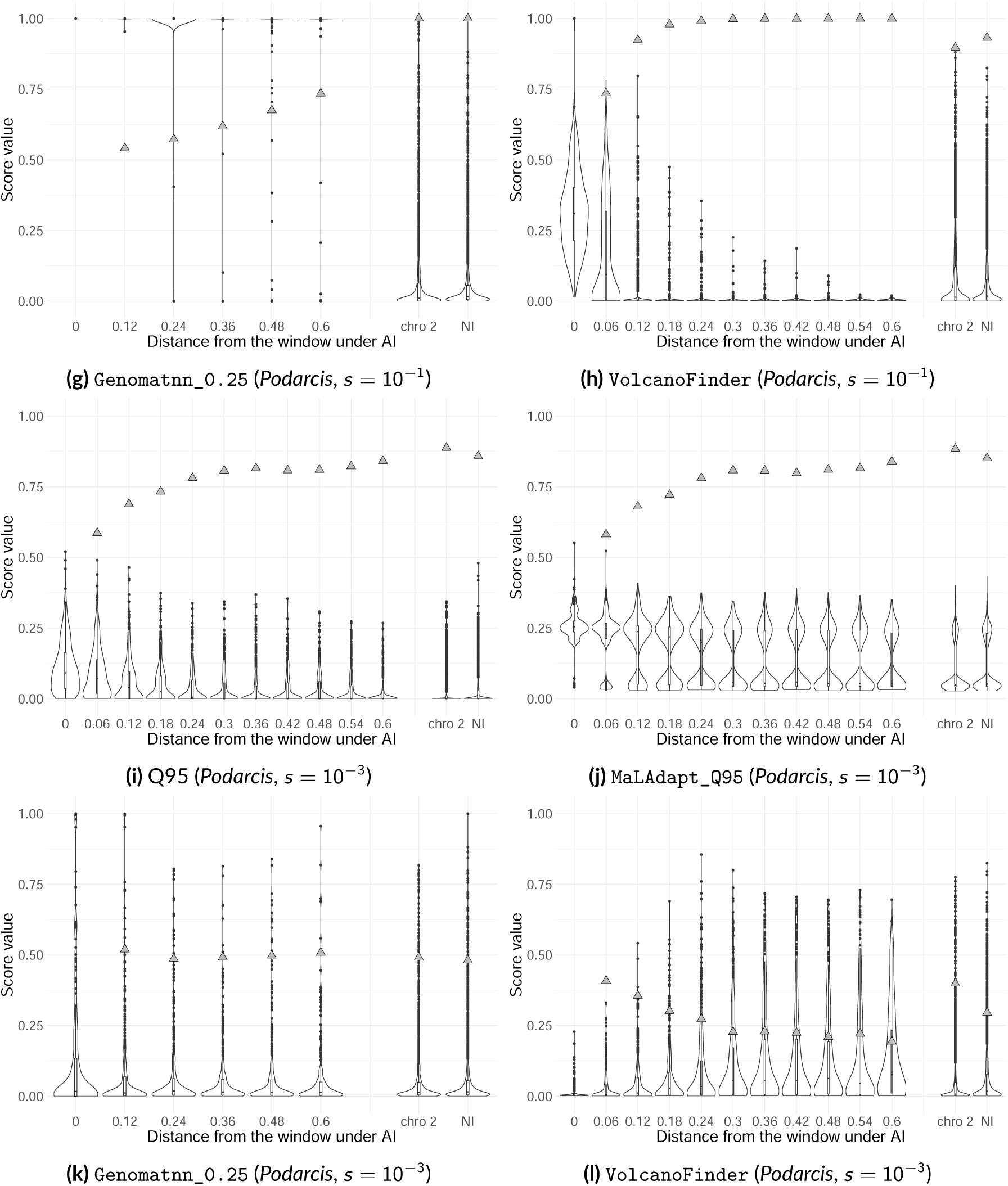
Score values of 200 test sets for the different windows considered under a *Podarcis* scenario with variable selection strength (intermediate, strong and weak). Scores from the first chromosome are represented as function of the distance (cM) of the windows to the site under selection (0 indicates the window containing it), followed by scores from the second neutral chromosome (“chro 2”) and from an independently simulated neutral chromosome (NI) for reference. The areas under the ROC curves (AUC) for each test set containing AI windows and each class of non-AI windows are represented by the grey triangles. Each violin plot is from at least 200 windows.

### Human reference scenario

#### ROC curves

The ROC curves in Fig. 2a shows the comparison of the performance of the methods under the human reference scenario when the non-AI windows are adjacent windows. Q95 (AUC = 0.978) appears to be the method with the best performance, closely followed by MaLAdapt_Q95 (AUC = 0.975), MaLAdapt_noQ95 (AUC = 0.968). Genomatnn_0.25 (AUC = 0.921) and Genomatnn_0.05 (AUC = 0.899) show poorer results and VolcanoFinder (AUC = 0.641) performances are much worse than all others.

All methods, except VolcanoFinder, perform better when non-AI windows are independent neutral introgressed (NI) windows (Fig. 3a and Table. S4) rather than adjacent windows. MaL-Adapt_Q95 shows a performance comparable to that of the Q95 (AUC = 0, 996 for both methods). The performance of MaLAdapt_noQ95 varies in the same way as MaLAdapt_Q95 but with the same or poorer AUC values, for most of the scenarios considered. The ranking of all methods remains the same as for adjacent windows (Q95 equal to MaLAdapt then Genomatnn followed by VolcanoFinder). In addition, the differences in AUC between the methods are much smaller than for the test sets with adjacent windows (except for VolcanoFinder).

Finally, considering the second neutral chromosome as non-AI windows (Fig. S3) does not change the relative performance of the methods, which is similar to classification performance against the NI windows.

#### Score distribution among window classes

Fig. 4 shows the variation of the score values of the different methods as a function of the distance to the window under AI. The expected pattern of decrease of the score with the distance, due to decreasing hitchhiking, is found for all methods except VolcanoFinder. Accordingly, AUC values increase with distance from the AI window for those methods. Both score and AUC variation patterns varies depending on the methods considered, showing much more spread out distributions of the score for Genomatnn than Q95 and MaLAdapt(Fig. 4), with many large score values present at larger distances. This results in a slower increase of the AUC for Genomatnn. On the opposite, VolcanoFinder’s scores are relatively constant for all distances, with only a slight increase at the window under AI (Fig. 4d).

As expected, due to intra-chromosomal hitchhiking, the scores of adjacent windows are distinct from those of NI windows, with higher mean scores even for the most distant windows. Conversely, inter-chromosomal hitchhiking seems to only have almost no effect, as the distributions of scores for the second chromosome windows are extremely close to those for the NI windows.

### Old divergence has a strong effect on performance (*Podarcis*)

#### ROC curves

When the divergence between the donor and recipient populations is much older than that of the human lineage (*i.e. Podarcis* scenario with intermediate selection), the performance of the methods increases, except for Genomatnn_0.05 and VolcanoFinder (Fig. 2b, Fig. 3b and Fig. S3b). Relative method performances stay the same, but MaLAdapt and Q95 sometimes show similar performance. With chro2 or NI windows as non-AI windows, both reach the AUC values of a perfect classifier (AUC = 1).

#### Score distribution among window classes

Under such old divergence, the scores also decrease with distance from the AI window due to hitchhiking, but only a little faster than under the human reference scenario. Accordingly, all methods reach AUC values close to 1 for shorter distances, except, as before, for VolcanoFinder (Fig. 5d) and Genomatnn_0.05 (Fig. S4d). The pattern differences between methods are similar to the previous cases, but Genomatnn (Fig. 5c) show a stronger score decline than under a more recent divergence, with very high scores around the introgressed mutation followed by an abrupt fall around 0.36 cM. At short distances, Q95 performs better than all other methods.

VolcanoFinder shows opposite patterns of variation with an increase of scores with distance from the AI window (Fig. 5d, but also performs extremely poorly under this scenario as previously shown by the ROC curves.

Finally, score values for the NI and chro2 windows scores are lower than for a more recent divergence but also lower than the scores of the most distant windows (0.6 cM) to the AI window on the first chromosome.

### Strength of selection has a strong effect on performance (*Podarcis*)

#### ROC curves

Under the two *Podarcis* scenarios with a stronger and a weaker selection strength (Fig. 2c and Fig. 2d), most methods perform less well and the order of method performance changed a little compared to the reference human scenario when the non-AI windows are adjacent windows. A moderate to strong decrease in performance is observed for all methods under a strong selection strength, except for VolcanoFinder for which performance is better than under any other scenario considered in this study (AUC = 0.961). In this situation, VolcanoFinder even outperforms Q95 (AUC = 0.944). Contrarily, Genomatnn_0.05 performs very poorly under strong selection (AUC = 0.051) and Genomatnn_0.25 is not better than a random classifier (AUC = 0.5). Under a weak selection strength, the drop in performance is even more important for all methods, with a best AUC = 0.76 for Q95. Interestingly, this is the only scenario where Genomatnn_0.05 outperforms Genomatnn_0.25, all other methods showing the same performance order than for intermediate selection strength (and the reference human scenario).

In contrast, when NI and chro2 windows are used as non-AI windows, all the methods perform better when selection is strong than in the presence of intermediate selection. All Q95, MaLAdapt and Genomatnn_0.25 methods reached AUC values equal to one. For weak selection, however, ROC curves for all methods show poorer performance than for intermediate selection, almost identical to the case of weak selection with adjacent windows as non-AI windows.

#### Score distribution among window classes

Under strong selection, a stronger hitchhiking effect increases the score values of all adjacent windows compared to intermediate selection for Q95, MaLAdapt and Genomatnn. The decay of the scores with distance to the AI site is thus much weaker and remains observable only for Q95 (Fig. 5e) and MaLAdapt (Fig. 5f). This stronger effect of hitchhiking thus increases the distance required for the methods to differentiate between AI and non-AI windows. For Genomatnn (Fig. 5g), both the AI window and adjacent one show large score values, greatly reducing its performances compared to intermediate selection (with only a very small increase of AUCs with distance and a low maximum value of AUC = 0.73). As seen for the ROC curves and unlike all other scenarios, VolcanoFinder is the best performing method under this strong selection and strong divergence scenario, showing perfect discrimination (AUC value of 1) for shorter distances to the AI window than Q95 (0.48 cM versus 0.54 cM).

Under strong selection, the scores of NI windows are much lower than those of the window under AI and adjacent ones for all methods, except VolcanoFinder, resulting in a perfect classification of the AI/non-AI windows for Q95, MaLAdapt and Genomatnn_0.25 (Fig. 5).

Inter-chromosomal hitchhiking is observed under strong selection and leads to larger scores for windows on the second chromosome than for NI windows for all methods. This effect is minimal for Genomatnn_0.25 (Fig. 5g) but stronger for Genomatnn_0.05, (Fig. S4f), with the second chromosome harbouring high score values typical of AI windows. Those high chro2 scores greatly reduce the performance of Genomatnn_0.05 when chro2 windows are used as non-AI, compared to NI windows (AUC = 0.66 against 0.90, respectively). A reduction of performance is also observed for VolcanoFinder with chro2 windows, but to a much lesser extent (AUC = 0.93 against 0.90, respectively). For MaLAdapt and Q95, the scores on the second chromosome are also higher than the scores for NI windows but still much lower than AI window scores (Fig. 5e and Fig. 5f), resulting in good performances.

When selection is weak (Fig. 5 and Fig. S4), scores of the AI window are much lower and the signal decay with distance is barely visible for Q95, MaLAdapt and Genomatnn_0.05 except for the closest adjacent windows (Fig. 5i, Fig. 5j and Fig. S4h), showing that the hitchhiking effect is very weak but present. For Q95, MaLAdapt and Genomatnn_0.05, a weak selection thus leads to an increase of AUC values with the distance from the AI site, but the AUCs never reach 1 for adjacent windows (maximum value reached by Q95: AUC = 0.84). For VolcanoFinder (Fig. 5l), the pattern of inverted score values previously observed for the *Podarcis* scenario with intermediate selection is still present, leading to very low AUC values. For Genomatnn_0.25, the distributions of scores for the AI windows and adjacent windows are almost identical, also resulting in low AUC values.

Scores of NI and chro2 windows are relatively similar due to the limited hitchhiking effect under weak selection, and thus lead to similar performances of all methods compared to results on adjacent windows.

### Old migration moderately reduces performance (*Ursus*)

#### ROC curves

The scenario with an old migration time (*Ursus*, *T_m_* = 15, 000 generations, Fig. 2e) leads to a reduction in performance for all the methods and all types of non-AI windows compared to the human reference scenario. This is the only case where the Q95 is not one of the best performing methods, outperformed by MaLAdapt and Genomatnn.

#### Score distribution among window classes

The AI window show lower scores for MaLAdapt and Q95 with this older migration event (Fig. S5a and Fig. S5e) than those with a more recent migration (*i.e.* the reference human scenario and the *Podarcis* one with the same intermediate selection strength). Adjacent windows have much more spread out distributions than those observed for the reference scenario for both methods. For Genomatnn the opposite effects are observed (Fig. S5d), with distribution spread out for the AI window, and reduced towards low values for adjacent windows.

With an old migration event, NI and chro2 windows show score patterns similar to those of the most distant adjacent windows for Q95, MaLAdapt, and Genomatnn.

For all AI and non-AI windows types, the distributions of score values are similar for VolcanoFinder (Fig. S5f), showing even less difference between AI and non-AI windows than observed for the reference scenario.

### Small effective population sizes strongly reduces performance (human)

#### ROC curves

ROC curves for the human scenario with small population sizes show the poorest performance for almost all methods among all the scenarios considered in this study (Fig. 2f). Q95, MaLAdapt and Genomatnn show similar poor performances, with all AUC values less than 0.75 for adjacent windows. Those poor performances are also found with NI windows Fig. S2e) but chro2 leads to better performances (except for VolcanoFinder, (Fig. S3f)). With chro2 windows as non-AI windows, the performance of all methods is only slightly lower than in the reference scenario (*e.g.* AUC = 0.988 for this scenario compared to 0.996 for the reference scenario, for Q95). For this scenario, based on AUC values, Genomatnn_0.05 becomes one of the best methods with Q95, outperforming MaLAdapt with NI and chro2 windows as non-AI windows.

#### Score distribution among window classes

Bimodal score distributions, without intermediate values, are observed for Q95 and MaL-Adapt for adjacent windows when population sizes are small (Fig. S6). For Genomatnn, adjacent windows show higher values than those under the reference human scenario (Fig. S6d).

Under such small population sizes, NI windows behave like adjacent windows for all methods. On the contrary, window scores from chro2 contrast completely with those for adjacent and NI windows and are lower than those of NI and adjacent windows, for all the methods. This is the scenario where the chro2 and NI windows have the most distinct score distributions.

As for the old migration scenario, the distributions of score values are similar for all AI and non-AI windows types for VolcanoFinder when population sizes are small (Fig. S6f).

### Migration rate has moderate effect on performance (human)

#### ROC curves

When the migration rate is high (human scenario with *m* = 10*^−^*^1^, Fig. S1a), a slight increase in performance is observed for Q95, MaLAdapt and VolcanoFinder with adjacent windows as non-AI windows compared to low and intermediate migration levels. On the opposite, a slight improvement is observed for Genomatnn when the migration rate is low (human scenario with *m* = 10*^−^*^3^, Fig. S1b) compared to high and intermediate migration levels. With both NI and chro2 windows as non-AI windows, Q95 and MaLAdapt are also very close to a perfect classifier (AUC = 0.999) for both low and high migrations rates, whereas Genomatnn’s performance remains slightly better with intermediate and low migration rates than with high migration.

#### Score distribution among window classes

Increasing the migration rate tends to shift the score distribution of all non-AI windows towards higher values, especially for Genomatnn (Fig. S7). For Q95 and MaLAdapt AI window scores also increase (Fig. S7e and Fig. S7a), but not for Genomatnn (Fig. S7d). Contrarily, a small migration rate (Fig. S8) leads to a decrease in both scores of the adjacent and AI windows for Q95 and MaLAdapt (Fig. S8c and Fig. S8a). For Genomatnn, only the AI windows have slightly higher scores and adjacent window scores are similar to those of the human reference scenario.

In the case of strong migration, NI and chro2 windows show higher scores than in the reference scenario, but with lower mean values than those of adjacent windows. For low migration, the NI and chro2 windows have score values similar to those observed for the reference scenario.

### Hotspots has a little effect on the performance (human)

#### ROC curves

The presence of a recombination hotspot at 20kb from the beneficial mutation tends to slightly reduce the performance of all methods when adjacent windows are used as non-AI windows (Fig. S1c), except Genomatnn whose performance increases slightly (AUC = 0.933 against 0.921 with and without hotspots, respectively). Conversely, with NI and chro2 windows as non-AI windows (Fig. S2c and S3i), the AUC of all methods except VolcanoFinder increases only by a few thousandths.

When the hotspot is on the advantageous mutation (Fig. S1d), all performance drops slightly for all types of non-AI window (*e.g.* AUC = 0.966 against 0.978 with and without the hotspot, respectively, for Q95).

#### Score distribution among window classes

The presence of a recombination hotspot distant by 20kb from the beneficial mutation has no major impact on the distribution of scores in the AI and non-AI windows (Fig. S9). Only adjacent windows close to the window under AI tend to have higher maximum values. This pattern is also present and stronger when the hotspot is in the AI mutation zone. Under this latter scenario, AI window scores also tend to decrease (Fig. S10). Hotspots have no effect on the second chromosome window scores and only a little effect on NI windows, apart from a small increase in the maximum values, causing the distributions of the NI windows to be slightly higher than those of chro2 for both hotspot scenarios.

### Balanced sample size slightly improves performance (human)

#### ROC curves

For all types of non-AI windows, balanced sample sizes, with more individuals from the donor populations and fewer individuals from the recipient ones compared to the reference human samples sizes, improve the performance of Q95, MaLAdapt, VolcanoFinder and Genomatnn_0.25 but decrease that of Genomatnn_0.05 (Fig. S1e, S2e and S3k).

#### Score distribution among window classes

Balanced sample sizes do not affect much the score distributions for the different window types, except a slight decrease of the scores of adjacent windows away from the beneficial mutation for Q95, MaLAdapt and Genomatnn_0.25 and slightly higher score values for AI windows for all methods (Fig. S11). With balanced sample sizes, MaLAdapt’s and Q95’s performances are indistinguishable for all types of non-AI windows considered.

## Discussion

Given our results, the use of Q95 as a genome scan statistic seems the most discriminating method, with high power for low false positive rates. It also leads to the narrowest regions of adjacent windows giving positive scores due to hitchhiking, allowing to better pinpoint the window under selection. Q95 is the best performing method in all but one of the scenario considered in this study, even under the weakest strength of selection or the small population size cases, the two scenarios showing the worst performances for all methods. The only scenario for which Q95 is not the most discriminating method is when the migration event is ancient (*i.e.* the *Ursus* scenario with the migration event at 15, 000 generations in the past). With such a long time after migration, genetic drift will lead to the fixation of random neutral introgressed tracts generating large Q95 values and, therefore, increasing false positive rates (FPR) on all non-AI window classes. To the extent that those results are robust, the Q95 summary statistic also appears quite convenient as it is easy to compute without training a machine learning algorithm, in contrast to simulation-based classification methods. Moreover, it does not require the use of phased data, unlike MaLAdapt, for example. Finally, Q95 also showed similar or even better performance than the other tested methods in previous comparisons (Gower et al., 2021; Racimo et al., 2017).

VolcanoFinder can be very powerful to detect AI when the mutation under AI has recently reached fixation and when the divergence between populations D and R is ancient (Gower et al., 2021; Setter et al., 2020; Zhang et al., 2023). For our human scenarios, the mutation under AI is generally not fixed and the divergence time is relatively recent. For the *Ursus* scenario (ancient migration), the mutation is indeed fixed but it is far too old for the AI patterns sought by Volcano-Finder to have persisted in the sequences. Only our *Podarcis* scenario with strong selection corresponds to a scenario where VolcanoFinder can have good performance (old divergence time, strong selection, recent migration and fixation of the mutation under AI). In previous evaluations on scenarios departing from these characteristics, VolcanoFinder was also shown to perform poorly compared to the other methods Gower et al. (2021) and Zhang et al. (2023). The presence of spurious peaks of VolcanoFinder’s scores on neutral regions near chromosome ends, leading to an increase of FPR, was also described in Setter et al. (2020).

Between the two simulation-based methods, MaLAdapt in general does better than Genomatnn, with a lower rate of false positives, especially in the presence of hitchhiking, in view of our results and those of Zhang et al. (2023). This result might have been expected, as adjacent windows are included in MaLAdapt training set but not in Genomatnn training. However, MaLAdapt also shows high FPR for the adjacent windows closest to the mutation from the AI windows.

The main difference between our results and previous works is a better performance reported by Zhang et al. (2023) for MaLAdapt and by Gower et al. (2021) for Genomatnn compared to Q95 in some of their scenarios, for which, contrarily to most of our test scenarios, all parameter values were included in their training set. The second difference that can be noted between our results and those of Zhang et al. (2023) concerns the effect of misspecification of the migration rate. In their results at low migration (5 × 10*^−^*^3^) the performance of MaLAdapt increased, but in our tests at low migration (*m* = 10*^−^*^3^) the performance decreased slightly compared to cases of higher migration (*m* = 10*^−^*^2^, *m* = 10*^−^*^1^). Neither of the low migration rates is included in the training set of MaLAdapt (training values being {0.01, 0.02, 0.05, 0.1}), so the slight drop in MaLAdapt performance in our simulations could be due to stronger misspecification.

Indeed, the performance of the two simulation-based classification methods, MaLAdapt and Genomatnn, necessarily depends on the training set. The differences in performance observed between our tests and those in the articles describing these methods can thus be partly explained by the fact that we used the pre-trained ML models of the original publications, rather than ML models specifically re-trained under our scenarios. Our test sets generated under various evolutionary scenarios thus corresponds to a case of misspecification of the MaLAdapt and Genomatnn’s models. Gower et al. (2021) showed that the performance of Genomatnn decreases when using test sets from a different demographic model than the one considered for training, and Q95 performed better than their method in this case. MaLAdapt’s performance also dropped when the simulated test data were produced under parameter values of the Genomatnn train set. Comparison between our results and their results is difficult because Zhang et al. (2023) averaged the performance of the methods (*i.e.* the false positive rates) over different scenarios, making it impossible to tease apart the effect of different factors (strength of selection, hitchhiking, etc.). In addition, it is also difficult to identify the exact reason behind this discrepancy from our results because the models used by Zhang et al. (2023) to generate test sets differ substantially from ours. Indeed, their model includes past changes in population size, occurrence of deleterious mutations and gene structure (intro/exon). Another possible cause, presented in Zhang et al. (2023) as a source of increased FPRs, is that MaLAdapt was trained on lower selection values than that corresponding to our strong selection scenario (*s* ∈ [10*^−^*^4^, 10*^−^*^2^] and *s* ∈ [10*^−^*^3^, 10*^−^*^1^] for MaLAdapt and our tests respectively), which might potentially explain the observed drop in its performance for non-AI adjacent windows with *s* = 10*^−^*^1^.

As discussed above, it is sometimes difficult to identify whether variations in a parameter have a real impact on AI signals when the parameter values chosen are in a case of model misspecification of a simulation-based inference method. To overcome the problem of misspecification, it is often advisable to re-train the methods with training data from a demographic scenario close to the biological model under study, or with wide ranges of parameter values when the latter is poorly known. However, this step of generating training sets is time-consuming and requires a great deal of computing power. In addition, the current versions of MaLAdapt and Genomatnn do not provide user-friendly tools for producing the training data under user-specified demographic scenarios. Therefore, application of these methods under an appropriate demographic model requires high investment by the user.

Our results show that false positive rates increase when considering non-AI window types not taken into account in the train set. The original train set of Genomatnn considers only independent neutral introgressed windows. Accordingly, our results on independent NI windows show coherent low false positive rates, at levels similar or only slightly higher than for Q95, but considering adjacent windows leads to higher false positive rate. Unlike Genomatnn, MaLAdapt took into account adjacent windows in its train set. This may explain, in part, why MaLAdapt performs better than Genomatnn whichever the type of non-AI window considered.

Our results highlight the importance of considering different types of neutral windows during training. The aim of all those classification methods is to specifically identify the window carrying the mutation under AI. It is therefore necessary to take into account windows that have undergone evolutionary processes that could lead to genetic signals similar to those of adaptive introgression. Independent neutral introgression is taken into account in train sets, but this processes is not the only ones that can leave confounding signals with AI in genomes. For example, classic selective sweep might also leave signals in the data similar to those of adaptive introgression. However, these methods have been shown to be generally robust to this form of selection (Gower et al., 2021; Racimo et al., 2017; Zhang et al., 2023), except VolcanoFinder which seems to have a moderate probability of classifying as AI the windows presenting strong selective sweeps (Setter et al., 2020). Finally, depending on the strength of the selection applied to the AI mutation and the number of migrants, the effect of hitchhiking can leave an AI signal on adjacent windows linked to the mutation. As seen in our results, the effect of hitchhiking can even extend, to a lesser extent, to other chromosomes, *i.e.* not carrying the mutation under AI. Our results thus also highlight the importance of taking into account intraand interchromosomal hitchhiking in the train set.

The parameters that have the greatest impact on the performance of the methods in our test sets appear to be the strength of the selection (*s*), the divergence times between the donor and recipient populations (*T_D_*_2_), the migration time (*T_m_*) and the population sizes (*N*). This conclusion is limited by the necessarily finite number of scenarios explored in this work and other processes could influence the performance of the methods. While we have not explored the performance of the methods under other demographic scenarios, such as bottlenecked populations or bidirectional introgression, our simulations provide some insight through the observed effects of genetic drift and differentiation between D and R. Another factor that has not been included in the published methods nor in our tests is spatial population structure, which could lead to false introgression signals in the genome due to incomplete lineage sorting (Tournebize and Chikhi, 2023). It is therefore important to take these confounding processes into account in the train set or during robustness tests in order to quantify and possibly limit their impact on the classification performances.

The use of classification methods implies the need to define a threshold for classification scores. This threshold can be defined for a given train set but can be difficult to define when trying to control the error rate when the proportions of positive cases in the analysed set differ from those in the train set. In this respect, binary classification problems for multiple windows in a genome are comparable to multiple hypothesis testing problems. The hypothesis tested here for each window is that there is no adaptive introgression within it. For genomic scan on multiple windows, it may be more specifically relevant to control the false discovery rate (FDR), *i.e.* the proportion of false positives among all positives detected.

Various strategies have been developed to control the FDR, such as methods based on the availability of p-values for each window (Benjamini and Hochberg, 1995), or empirical Bayes procedures requiring the distribution of some “score” for each window under the null hypothesis (Efron, 2010; Storey, 2003). All these methods require the distribution of classification scores (or p-values) under the null model to be known, so it may be necessary to run simulations under the null model, even for scoring methods that do not require training (such as Q95). Some of the methods to control the FDR require independence among windows scores, which is not the case for our inferences because the presence of AI windows affects the distribution of scores in adjacent non-AI windows through hitchhiking. For this reason, an FDR method that takes data dependency into account would be required (Heller and Rosset, 2021; Tang and Zhang, 2007). Here, it might be further necessary to control the error rate among non-AI windows even in the presence of some AI window(s) in a genome, which may raise distinct problems in controlling error rates.

For such reasons, it might be necessary to first infer a single measure of AI at the genomic level (rather that at the level of individual windows), such as an overall adaptive effect of AI representing a cumulative effect of adaptive mutations in different windows, and then to use windows-based criteria measures of adaptive effect to identify the windows likely to contribute to the genomic level measure of AI. It is possibly in this context that the windows-based methods evaluated in this paper could be best used.

## Acknowledgements

We would like to thank Graham Gower for his availability, help and advice in using Genomatnn. We would also like to thank Xinjun Zhang for giving us permission to use her scripts and for answering all our questions about MaLAdapt. Finally, we would also like to thank Sylvain Mousset for his help and advice on using VolcanoFinder.

We would also like to thank the recommander, Amandine Cornille and all the reviewers, Thibault Leroy, Lukas Metzger and Fernando Racimo whose suggestions and comments helped improve this manuscript.

## Data, script, code, and supplementary information availability

Scripts and code are available online (https://doi.org/10.5281/zenodo.14181497; Romieu et al., 2024a).

Supplementary information is available online (https://doi.org/10.5281/zenodo.14205482; Romieu et al., 2024b).

## References

Adavoudi R, Pilot M (2022). Consequences of hybridization in mammals: a systematic review. Genes 13. 50. 10.3390/genes13010050.

Adrion JR, Cole CB, Dukler N, Galloway JG, Gladstein AL, Gower G, Kyriazis CC, Ragsdale AP, Tsambos G, Baumdicker F, Carlson J, Cartwright RA, Durvasula A, Gronau I, Kim BY, McKenzie P, Messer PW, Noskova E, Ortega-Del Vecchyo D, Racimo F, et al. (2020). A community-maintained standard library of population genetic models. eLife 9, e54967. 10.7554/eLife.54967.

Anderson E, Hubricht L (1938). Hybridization in Tradescantia. III. The evidence for introgressive hybridization. American Journal of Botany 25. 396–402. 10.1002/j.1537-2197.1938.tb09237.x.

Baumdicker F, Bisschop G, Goldstein D, Gower G, Ragsdale AP, Tsambos G, Zhu S, Eldon B, Ellerman EC, Galloway JG, Gladstein AL, Gorjanc G, Guo B, Jeffery B, Kretzschumar WW, Lohse K, Matschiner M, Nelson D, Pope NS, Quinto-Cortés CD, et al. (2022). Efficient ancestry and mutation simulation with msprime 1.0. Genetics 220, iyab229. 10.1093/genetics/iyab229.

Benjamini Y, Hochberg Y (1995). Controlling the false discovery rate: a practical and powerful approach to multiple testing. Journal of the Royal Statistical Society: Series B (Methodological) 57. 289–300. 10.1111/j.2517-6161.1995.tb02031.x.

Bradley AP (1997). The use of the area under the ROC curve in the evaluation of machine learning algorithms. Pattern Recognition 30, 1145–1159. 10.1016/S0031-3203(96)00142-2.

Burgarella C, Barnaud A, Kane NA, Jankowski F, Scarcelli N, Billot C, Vigouroux Y, Berthouly-Salazar C (2019). Adaptive introgression: an untapped evolutionary mechanism for crop adaptation. Frontiers in Plant Science 10. 4. 10.3389/fpls.2019.00004.

Burke JM, Arnold ML (2001). Genetics and the fitness of hybrids. Annual Review of Genetics 35. 31–52. 10.1146/annurev.genet.35.102401.085719.

Caeiro-Dias G, Brelsford A, Kaliontzopoulou A, Meneses-Ribeiro M, Crochet PA, Pinho C (2021). Variable levels of introgression between the endangered *Podarcis carbonelli* and highly divergent congeneric species. Heredity 126. 463–476. 10.1038/s41437-020-00386-6.

Caeiro-Dias G, Brelsford A, Meneses-Ribeiro M, Crochet PA, Pinho C (2023). Hybridization in late stages of speciation: Strong but incomplete genome-wide reproductive isolation and ‘large Z-effect’ in a moving hybrid zone. Molecular Ecology 32. 4362–4380. 10.1111/mec.17035.

Dabi A, Schrider DR (2024). Population size rescaling significantly biases outcomes of forwardin-time population genetic simulations. *Genetics*, iyae180. 10.1093/genetics/iyae180.

Edelman NB, Mallet J (2021). Prevalence and adaptive impact of introgression. Annual Review of Genetics 55. 265–283. 10.1146/annurev-genet-021821-020805.

Efron B (2010). Large-scale inference: empirical Bayes methods for estimation, testing, and prediction. In: Institute of Mathematical Statistics monographs. Cambridge University Press, p. 276. 10.1017/CBO9780511761362.

Gaczorek TS, Chechetkin M, Dudek K, Caeiro-Dias G, Crochet PA, Geniez P, Pinho C, Babik W (2023). Widespread introgression of MHC genes in Iberian *Podarcis* lizards. Molecular Ecology 32. 4003–4017. 10.1111/mec.16974.

Geurts P, Ernst D, Wehenkel L (2006). Extremely randomized trees. Machine Learning 63, 3–42. 10.1007/s10994-006-6226-1.

Gower G, Picazo PI, Fumagalli M, Racimo F (2021). Detecting adaptive introgression in human evolution using convolutional neural networks. eLife 10. e64669. 10.7554/eLife.64669.

Grant PR, Grant BR (2019). Hybridization increases population variation during adaptive radiation. Proceedings of the National Academy of Sciences 116. 23216–23224. 10.1073/pnas.1913534116.

Haller BC, Messer PW (2019). SLiM 3: Forward genetic simulations beyond the Wright–Fisher model. Molecular Biology and Evolution 36, 632–637. 10.1093/molbev/msy228.

Harrison RG, Larson EL (2014). Hybridization, introgression, and the nature of species boundaries. Journal of Heredity 105, 795–809. 10.1093/jhered/esu033.

Hedrick PW (2013). Adaptive introgression in animals: examples and comparison to new mutation and standing variation as sources of adaptive variation. Molecular Ecology 22. 4606– 4618. 10.1111/mec.12415.

Heller R, Rosset S (2021). Optimal control of false discovery criteria in the two-group model. Journal of the Royal Statistical Society Series B: Statistical Methodology 83, 133–155. 10.1111/rssb.12403.

Hoggart CJ, Chadeau-Hyam M, Clark TG, Lampariello R, Whittaker JC, De Iorio M, Balding DJ (2007). Sequence-level population simulations over large genomic regions. Genetics 177, 1725–1731. 10.1534/genetics.106.069088.

Jo BS, Choi SS (2015). Introns: The functional benefits of introns in genomes. Genomics & Informatics 13. 112–118. 10.5808/GI.2015.13.4.112.

Kaliontzopoulou A, Phino C, Harris DJ, Carretero MA (2011). When cryptic diversity blurs the picture: a cautionary tale from Iberian and North African *Podarcis* wall lizards. Biological Journal of the Linnean Society 103, 779–800. 10.1111/j.1095-8312.2011.01703.x.

Kelleher J, Etheridge AM, McVean G (2016). Efficient coalescent simulation and genealogical analysis for large sample sizes. PLOS Computational Biology 12. e1004842. 10.1371/journal.pcbi.1004842.

Kelly JK (1997). A test of neutrality based on interlocus associations. Genetics 146, 1197–1206. 10.1093/genetics/146.3.1197.

Kim SC, Rieseberg LH (1999). Genetic architecture of species differences in annual sunflowers: implications for adaptive trait introgression. Genetics 153, 965–977. 10.1093/genetics/153.2.965.

Lan T, Leppälä K, Tomlin C, Talbot SL, Sage GK, Farley SD, Shideler RT, Bachmann L, Wiig Ø, Albert VA, Salojärvi J, Mailund T, Drautz-Moses DI, Schuster SC, Herrera-Estrella L, Lindqvist C (2022). Insights into bear evolution from a Pleistocene polar bear genome. Proceedings of the National Academy of Sciences 119, e2200016119. 10.1073/pnas.2200016119.

Liu S, Lorenzen ED, Fumagalli M, Li B, Harris K, Xiong Z, Zhou L, Korneliussen TS, Somel M, Babbitt C, Wray G, Li J, He W, Wang Z, Fu W, Xiang X, Morgan CC, Doherty A, O’Connell MJ, McInerney JO, et al. (2014). Population genomics reveal recent speciation and rapid evolutionary adaptation in polar bears. Cell 157, 785–794. 10.1016/j.cell.2014.03.054.

Liu S, Zhang L, Sang Y, Lai Q, Zhang X, Jia C, Long Z, Wu J, Ma T, Mao K, Street NR, Ingvarsson PK, Liu J, Wang J (2022). Demographic history and natural selection shape patterns of deleterious mutation load and barriers to introgression across *Populus* genome. Molecular Biology and Evolution 39, msac008. 10.1093/molbev/msac008.

Mallet J (2005). Hybridization as an invasion of the genome. Trends in Ecology & Evolution 20, 229–237. 10.1016/j.tree.2005.02.010.

Maynard Smith J, Haigh J (1974). The hitch-hiking effect of a favourable gene. Genetics Research 23, 23–35. 10.1017/S0016672300014634.

Miles A, Rodrigues MF, Ralph P, Kelleher J, Schelker M, Pisupati R, Rae S, Millar T (2024). scikit-allel: v1.3.8. 10.5281/zenodo.10876220.

Myers S, Spencer C, Auton A, Bottolo L, Freeman C, Donnelly P, McVean G (2006). The distribution and causes of meiotic recombination in the human genome. Biochemical Society Transactions 34, 526–530. 10.1042/bst0340526.

Pardo-Diaz C, Salazar C, Baxter SW, Merot C, Figueiredo-Ready W, Joron M, McMillan WO, Jiggins CD (2012). Adaptive introgression across species boundaries in *Heliconius* butterflies. PLOS Genetics 8. e1002752. 10.1371/journal.pgen.1002752.

Pawar H, Rymbekova A, Cuadros-Espinoza S, Huang X, Manuel M, Valk T, Lobon I, Alvarez-Estape M, Haber M, Dolgova O, Han S, Esteller-Cucala P, Juan D, Ayub Q, Bautista R, Kelley JL, Cornejo OE, Lao O, Andrés AM, Guschanski K, et al. (2023). Ghost admixture in eastern gorillas. Nature Ecology & Evolution 7. 1503–1514. 10.1038/s41559-023-02145-2.

Pinho C, Kaliontzopoulou A, Carretero MA, Harris DJ, Ferrand N (2009). Genetic admixture between the Iberian endemic lizards *Podarcis bocagei* and *Podarcis carbonelli*: evidence for limited natural hybridization and a bimodal hybrid zone. Journal of Zoological Systematics and Evolutionary Research 47. 368–377. 10.1111/j.1439-0469.2009.00532.x.

Pinho C, Kaliontzopoulou A, Harris DJ, Ferrand N (2011). Recent evolutionary history of the Iberian endemic lizards *Podarcis bocagei* (Seoane, 1884) and *Podarcis carbonelli* Pérez-Mellado, 1981 (Squamata: Lacertidae) revealed by allozyme and microsatellite markers. Zoological Journal of the Linnean Society 162, 184–200. 10.1111/j.1096-3642.2010.00669.x.

Prüfer K, Racimo F, Patterson N, Jay F, Sankararaman S, Sawyer S, Heinze A, Renaud G, Sudmant PH, Filippo C, Li H, Mallick S, Dannemann M, Fu Q, Kircher M, Kuhlwilm M, Lachmann M, Meyer M, Ongyerth M, Siebauer M, et al. (2014). The complete genome sequence of a Neanderthal from the Altai Mountains. Nature 505. 43–49. 10.1038/nature12886.

Racimo F, Marnetto D, Huerta-Sánchez E (2017). Signatures of archaic adaptive introgression in present-day human populations. Molecular Biology and Evolution 34, 296–317. 10.1093/molbev/msw216.

Racimo F, Sankararaman S, Nielsen R, Huerta-Sánchez E (2015). Evidence for archaic adaptive introgression in humans. Nature Reviews Genetics 16. 359–371. 10.1038/nrg3936.

Ralph P, Thornton K, Kelleher J (2020). Efficiently summarizing relationships in large samples: A general duality between statistics of genealogies and genomes. Genetics 215, 779–797. 10.1534/genetics.120.303253.

Renoult JP, Geniez P, Bacquet P, Benoit L, Crochet PA (2009). Morphology and nuclear markers reveal extensive mitochondrial introgressions in the Iberian wall lizard species complex. Molecular Ecology 18. 4298–4315. 10.1111/j.1365-294X.2009.04351.x.

Romieu J, Camarata G, André-Crochet P, Navascués M, Leblois R, Rousset F (2024a). IntroAdapt-Methods: Scripts for “Performance evaluation of adaptive introgression classification methods”. 10.5281/zenodo.14181498.

Romieu J, Camarata G, Crochet PA, Navascués M, Leblois R, Rousset F (2024b). Supplementary Information for “Performance evaluation of adaptive introgression classification methods”. Version 2. 10.5281/zenodo.14205482.

Rosser N, Seixas F, Queste LM, Cama B, Mori-Pezo R, Kryvokhyzha D, Nelson M, Waite-Hudson R, Goringe M, Costa M, Elias M, Mendes Eleres de Figueiredo C, Freitas AVL, Joron M, Kozak K, Lamas G, Martins ARP, McMillan WO, Ready J, Rueda-Muñoz N, et al. (2024). Hybrid speciation driven by multilocus introgression of ecological traits. Nature 628. 811–817. 10.1038/s41586-024-07263-w.

Runemark A, Vallejo-Marin M, Meier JI (2019). Eukaryote hybrid genomes. PLOS Genetics 15. e1008404. 10.1371/journal.pgen.1008404.

Setter D, Mousset S, Cheng X, Nielsen R, DeGiorgio M, Hermisson J (2020). VolcanoFinder: Genomic scans for adaptive introgression. PLOS Genetics 16. e1008867. 10.1371/journal.pgen.1008867.

Storey JD (2003). The positive false discovery rate: a Bayesian interpretation and the q-value. The Annals of Statistics 31. 2013–2035. 10.1214/aos/1074290335.

Suarez-Gonzalez A, Lexer C, Cronk QCB (2018). Adaptive introgression: a plant perspective. Biology Letters 14. 20170688. 10.1098/rsbl.2017.0688.

Tang W, Zhang CH (2007). Empirical Bayes methods for controlling the false discovery rate with dependent data. In: Institute of Mathematical Statistics Lecture Notes - Monograph Series. Institute of Mathematical Statistics, pp. 151–160. 10.1214/074921707000000111.

Taylor SA, Larson EL (2019). Insights from genomes into the evolutionary importance and prevalence of hybridization in nature. Nature Ecology & Evolution 3. 170–177. 10.1038/s41559-018-0777-y.

Todesco M, Pascual MA, Owens GL, Ostevik KL, Moyers BT, Hübner S, Heredia SM, Hahn MA, Caseys C, Bock DG, Rieseberg LH (2016). Hybridization and extinction. Evolutionary Applications 9. 892–908. 10.1111/eva.12367.

Tournebize R, Chikhi L (2023). Questioning Neanderthal admixture: on models, robustness and consensus in human evolution. 10.1101/2023.04.05.535686. eprint: https://www.biorxiv.org/content/early/2023/04/05/2023.04.05.535686.full.pdf.

Twyford AD, Ennos RA (2012). Next-generation hybridization and introgression. Heredity 108. 179–189. 10.1038/hdy.2011.68.

Wang Y, Wang Y, Cheng X, Ding Y, Wang C, Merilä J, Guo B (2023). Prevalent introgression underlies convergent evolution in the diversification of *Pungitius* sticklebacks. Molecular Biology and Evolution 40, msad026. 10.1093/molbev/msad026.

Yang W, Feiner N, Pinho C, While GM, Kaliontzopoulou A, Harris DJ, Salvi D, Uller T (2021). Extensive introgression and mosaic genomes of Mediterranean endemic lizards. Nature Communications 12. 2762. 10.1038/s41467-021-22949-9.

Zhang X, Kim B, Singh A, Sankararaman S, Durvasula A, Lohmueller KE (2023). MaLAdapt reveals novel targets of adaptive introgression from Neanderthals and Denisovans in worldwide human populations. Molecular Biology and Evolution 40, msad001. 10.1093/molbev/msad001.

